# A gene co-expression network-based analysis of mesenchymal stromal cells reveals novel genes and molecular pathways underlying heterotopic ossification

**DOI:** 10.1101/2024.12.01.626232

**Authors:** Meiling Yang, Yongpeng Shi, Shichao Shen, Bin Yan, Lei Yang, Liuxun Li

**Affiliations:** Department of Oncology, Shenzhen Hospital of Guangzhou University of Chinese Medicine (Futian), Shenzhen, Guangdong, China; Department of spine surgery/Orthopaedics, the First Affiliated Hospital, Shenzhen University, Shenzhen Second People’s Hospital, Shenzhen, Guangdong, China

**Keywords:** Heterotopic ossification, Hub genes, Immune infiltration, Potential therapeutic agents, Weighted gene co-expression network analysis

## Abstract

**Background:** Heterotopic ossification (HO) represents a frequently seen refractory disease second to musculoskeletal injury, in which bone tissue ectopically exists within soft tissue, resulting in serious extremity loss-of-function. This work focused on identifying regulating factors and gene network associated with HO pathology.

**Material and Methods:** We randomized the heterotopic ossification dataset GSE94683 as HO and NON-HO group. Weighted gene co-expression network analysis (WGCNA) was applied in identifying HO related modules. We discovered differentially expressed genes (DEGs) between the two groups. Then, we integrated protein-protein interaction (PPI) network, co-expression network, enrichment analysis, gene set variation analysis (GSVA) and gene set enrichment analysis (GSEA) for identifying the HO-related pathways. We also employed GSE126118 test set to investigate hub genes related to HO. Eventually, potential therapeutics for reversing abnormal hub gene levels were predict by DGIdb.

**Results:** We discovered twelve HO status modules and 1,483 DEGs in HO samples compared with NON-HO counterparts. Four hub genes were obtained from the overlap of HO related PPI and coexpression networks. Training and test sets were adopted for verification, and three abnormally high hub gene expression in HO were obtained. Functional enrichment of DEGs indicated that these genes were involved in the stem cell pluripotency regulatory pathways, cytokine-cytokine receptor interaction, as well as rap1 pathway. Based on the results from GSEA and GSVA, many hub gene sets were mainly enriched in immune responses.

**Conclusion:** We identified a gene coexpression network associated with regulatory factors in HO, which provided a new perspective on the pathogenesis and provided potential biomarkers or therapeutic targets.

## Introduction

Heterotopic ossification (HO) accounts for a frequently occurring soft tissue disorder following injury [1]. It has the feature of pathological bone formation in the soft tissues resulting in joint swelling, pain, reduced range of motion and loss of functional capacity. The incidence of HO can be as high as 90% following acetabular fracture and total hip arthroplasty (THA) [2, 3]. However, its pathogenesis is not completely clear. HO has the same pathologies as additional frequently seen, refractory diseases like inflammatory response, immune response, resulting in perpetuates and recurrence. At present, surgery remains the mere efficient way to treat HO, however, it just has transient effect, because HO will recur, even 15 years postoperatively [4]. Additionally, surgical complications may take place in the case of ossification entrapping nerves and great blood vessels [5]. Therefore, it is necessary to identify regulators to shed more lights on designing efficient therapeutics.

Previous research has conducted transcriptomic analysis for reporting that dynamic alterations in hub genes and biological processes within the mouse burn/tenotomy-induced HO model and spinal cord injury-induced neurogenic heterotopic ossification model [6, 7]. Another recent study applied gene expression microarrays to identify HO related biomarkers and pathways [8]. Nevertheless, the approaches utilized in those works usually examine individual genes or co-expression genes that show close biological effects by gene functions in vivo. HO indicates a complicated pathological event possibly resulting from several genes. Consequently, simultaneous measurement of influence of different gene variants is helpful in determining the causative factor of disease. In addition, synergic effect induced by several gene changes is seen in the pathological processes of HO [9]. WGCNA can serve as a new tool to analyze the gene expression pattern for different specimens with specific clinical traits [10]. Unlike previously screened DEGs, WGCNA aggregates genes with close correlation to a module, later. it associates these genes with clinical features, and this helps to identify diagnostic biomarkers as well as therapeutic targets [11]. So far, WGCNA is widely used for transcriptomics studies (e.g. glioblastoma [12], Kawasaki disease [13], schizophrenia spectrum [14], etc.) Thus, it is speculated that the recognition of such co-expression modules can help to further elucidate the disease-related pathological process. Therefore, analyzing changing of co-expression gene network may be important in the pathological process of HO, so as to explore the molecular basis of this morphological change and the regulation of local microenvironment.

In this study, WGCNA and PPI networks were adopted to comprehensive analyze the HO related hub genes and biological pathways. Then, we employed the training and test sets for detecting hub gene expression, respectively. Later, we conducted Gene Ontology (GO) functional annotation together with Kyoto Encyclopedia of Genes and Genomes (KEGG), and Reactome pathway enrichment, for exploring possible underlying molecular mechanisms in HO. For exploring the pathogenesis of HO, GSVA and GSEA were employed by different samples as phenotype label. In addition, CIBERSORT convolution algorithm was performed for analyzing immune infiltrating degrees. For improving HO treatment, DGIdb database was carried out to predict potential drugs and molecular compounds. The above results are conductive to identifying new meaningful biomarkers and exploring HO pathogenesis, thus contributing to diagnosing and treating HO. Figure 1 exhibits the research flowchart.

**Figure 1.**
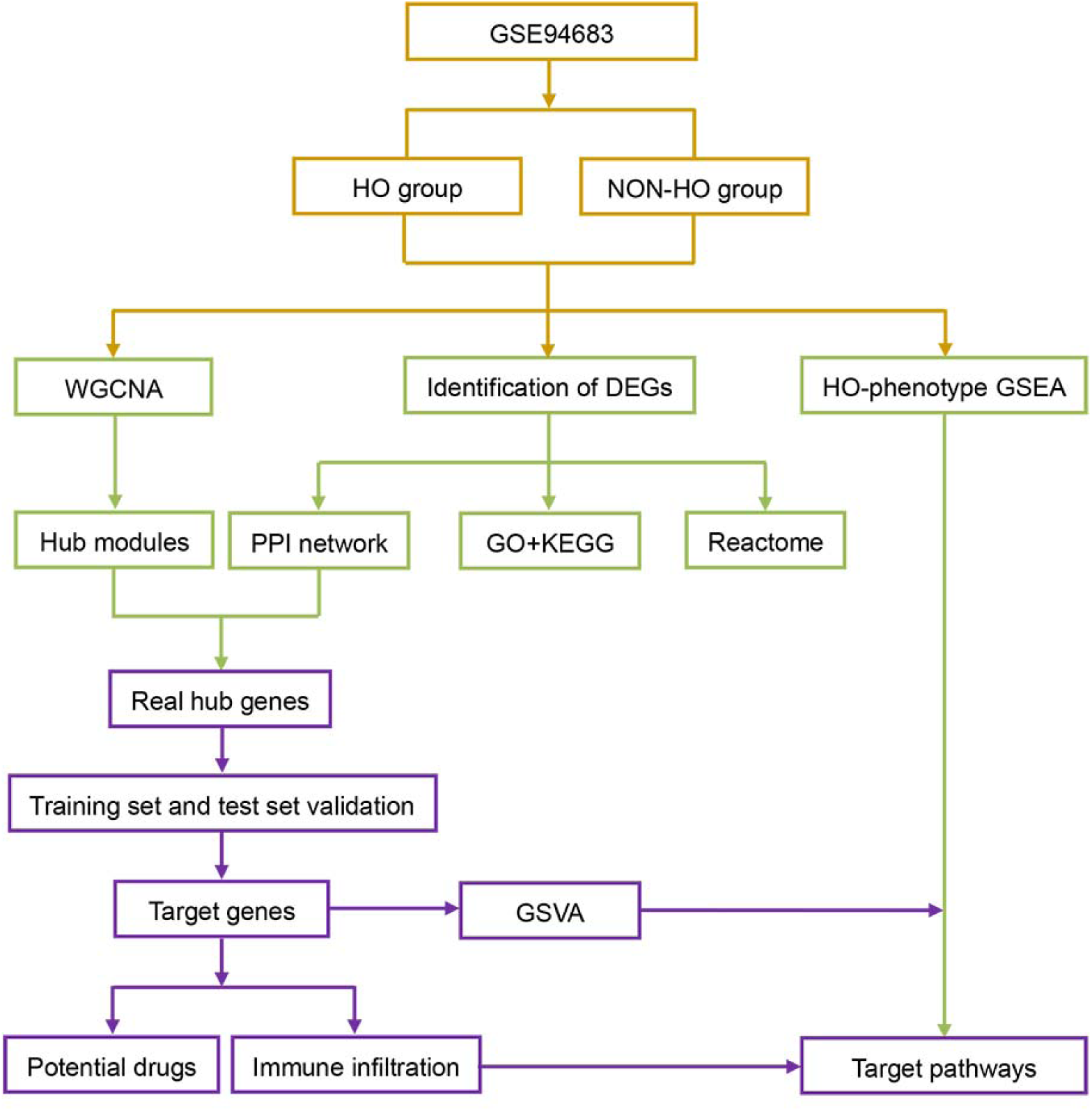
Research flowchart.

## Methods

### Search strategy

In this study, we obtained mRNA expression data of heterotopic ossification (HO) in GEO database from NCBI (http://www.ncbi.nlm.nih.gov/geo/) using the keywords “heterotopic ossification” until December 1^st^, 2024. Our retrieval strategy was designed below: (a) Gene expression profile data set using microarray chip technology; (b) Articles that compared gene expression profiles between HO samples and NON-HO samples; (c) The organism was homo sapiens. We removed articles that did not satisfy these criteria. Studies that do not meet the above criteria are excluded. The above search strategy was performed by two reviewers independently.

### Construction of HO-related co-expression network

To find HO highly relevant modules, WGCNA was employed through the WGCNA package in R language and carried out using the whole genes [15, 16]. Based on the connection strength adjacency matrix for gene pairs, an unsupervised co-expression network was established by Pearson correlation coefficient. The matrix was added to β = 7 on the basis of the scale-free topology (SFT) criterion. Then the adjacency matrix of clustering HO-related gene expression data was analyzed by topological overlap matrix. Last, a dynamic tree cutting algorithm was adopted to generate the dendrogram for module recognition. The minimum size of the number of module genes is set to 50, and the cutting height is 0.9.

### Identification of HO-related hub module

For identifying the modules markedly related to disease phenotype characteristics (HO vs. NON-HO), MEs (representing the first principal component (PC1) within the module) [17] was related to external characteristics for identifying most significant correlation. Module membership (MM) was defined as association of module eigengene with gene levels. Gene significance (GS) index refers to the absolute value relationship between gene and external traits. Meantime, we determined GS for measuring association of individual gene with external clinical trait. In the present study, genes showing the highest MM and GS values within our interested modules were recognized to be the natural candidate genes for subsequent analysis [18–21].

### Establishment of HO-related PPI network

HO related DEGs were imported into Search Tool for the Retrieval of Interacting Genes/Proteins (STRING) (version 1.4.0) (https://string-db.org/) [22] and confidence > 0.4 was select to exported to the Cytoscape (Version 3.9.0) [23] for constructing a PPI network. CytoHubba plugin was chosen to calculate and visualize the interactions between two proteins. The nodes and edges with higher degree of connection between proteins play vital roles in maintaining PPI network stability.

### Identification of HO-related hub genes between hub module and PPI network

Hub genes are intrinsically associated with additional genes in a module, and they have been considered to be functionally significant in previous studies. First, we screened the hub gene in HO phenotype-associated module co-expression network. Second, we screened the top 5% genes in PPI and co-expression networks as hub genes, respectively [24–26]. Third, we considered those sharing hub genes between PPI and co-expression networks as the “real” hub genes, which were chose for subsequent analyses. For hub genes validation, the datasets were divided into training sets and validation sets. As for GSE94683 training set and GSE126118 test set, HO was compared to the normal control group for true hub genes. *P <* 0.05 stood for statistical significance. GraphPad Prism (version, 8.0; GraphPad Software, Inc., La Jolla, Ca, USA) was employed for figure plotting.

### Functional annotations of HO-related hub gene

GO functional enrichment analysis has become an effective method for large-scale functional enrichment. KEGG has been the extensively adopted database, which preserves excessive data regarding diseases, genomes, biological pathways (BPs), drugs and chemical substances. In this study, we conducted GO as well as KEGG analysis on the DEGs using R package clusterProfiler [27]. In addition, Reactome knowledgebase (https://reactome.org/) [28–30] offers detailed molecular data for metabolism, DNA replications, signals, transportation, as well as additional cell events. In the current work, Reactome was utilized for identifying the top 10 BPs. *P* < 0.05 stood for statistical significance.

### Gene Set Variation Analysis (GSVA) and Gene set enrichment analysis (GSEA)

GSVA (version 1.22.0) [31] represents an approach to enrich gene sets, which estimates the changes in pathway activities in the samples. GSVA was carried out in this study for analyzing KEGG enrichment pathways on the basis of thresholds for enrichment score change > 1.0 and *P*-value < 0.05. Meantime, we conducted GSEA (version 4.1.0) [32] for investigating HO-related biological pathways. Enriched terms were conducted upon the *P* < 0.05 and false discovery rate (FDR) *q* < 0.25 thresholds.

### Prediction of HO-related immune infiltrating degrees

Different immune infiltrating degrees in HO compared with NON-HO groups within 16 infiltrating immune cells and 13 immune characteristics were carried out through CIBERSORT deconvolution algorithm [33]. We considered overlapping items showing an identical trend as alterations of immune characteristics. For investigating how HO immune infiltrating degrees affected 29 TIICs, the GSE94683 dataset was utilized for single-sample gene set enrichment analysis (ssGSEA) [31]. *P* < 0.05 stood for statistical significance.

### Potential drug identification

For identifying candidate drug targets for our discovered genes, this study employed Drug-Gene Interaction database (DGIdb) (http://dgidb.genome.wustl.edu/) for analysis. This database covered data associated with ‘druggable genes’, human drugs, as well as drug-gene interactions of 13 diverse sources; at present, it covers over 2,611 human genes, 6,307 drugs and 14,144 drug-gene interactions [34] . We conducted DGIdb for predicting possible drugs and molecular compounds that interacted with the hub genes. We also utilized Cytoscape software for visualizing drug–gene interaction network.

## Results

### Included study characteristics

After a careful screening, we obtained two microarray datasets (GSE94683 and GSE126118) in NCBI GEO database [7, 35–37]. GSE94683 dataset was obtained on the basis of the platform Agilent-021531 Whole Human Genome Oligo Microarray 4x44K, while GSE126118 on the basis of platform GPL13112 Illumina HiSeq 2000 (Mus musculus). We collected GSE94683 microarray data from mesenchymal stromal cells (MSCs) from NON-HO patients (HO group, n=7) and from bone marrow of healthy donors (NON-HO, n=9). The GSE126118 dataset covered samples collected in 7 groups, which included 2 from tenotomy site of a burn/tenotomy mouse (3 weeks after injury) strain, 3 from uninjured contralateral hindlimb tendon of a burn/tenotomy mouse (3 weeks after injury), and 2 from uninjured hindlimb tendon (no burn and no tenotomy). In the current work, GSE94683 dataset was used to establish PPI and co-expression networks for screen “real” hub genes as well as pathways related to HO. Meanwhile, GSE126118 was defined as the test set for validation of real hub genes.

### Construction of the co-expression network and identification of HO-related modules

Co-expression network construction and HO related modules were carried out via WGCNA using the R language “WGCNA” package. In this study, power β = 10 was chosen as soft-thresholding parameter for ensuring the scale-free network. Sixteen samples were observed to classify into 2 clusters. Additionally, correlation of modules was examined by Pearson’s correlation. We excavated altogether 12 modules, with the lightgreen module having the closest relationship with HO traits (Fig. 2A-B). According to these findings, each module is independently verified, which indicates that each module has a high degree of independence and the relative gene expression independence of each module. For exploring co-expression similarities among these 12 modules, first the connectivity of characteristic genes was evaluated, then the consensus correlation cluster analysis was carried out (Figure 2C). Furthermore, the 12 modules were respectively analyzed within modules, which included GS and MM. We calculated the lightgreen module for better exploring closely associated genes. Fig. 2D displays a scatter diagram of GS about HO characteristics, and the disease status relative to MM in light green module. HO, MM and GS showed significant positive correlations, indicating the possible relation between the most important lightgreen module with these external traits.

**Figure 2.**
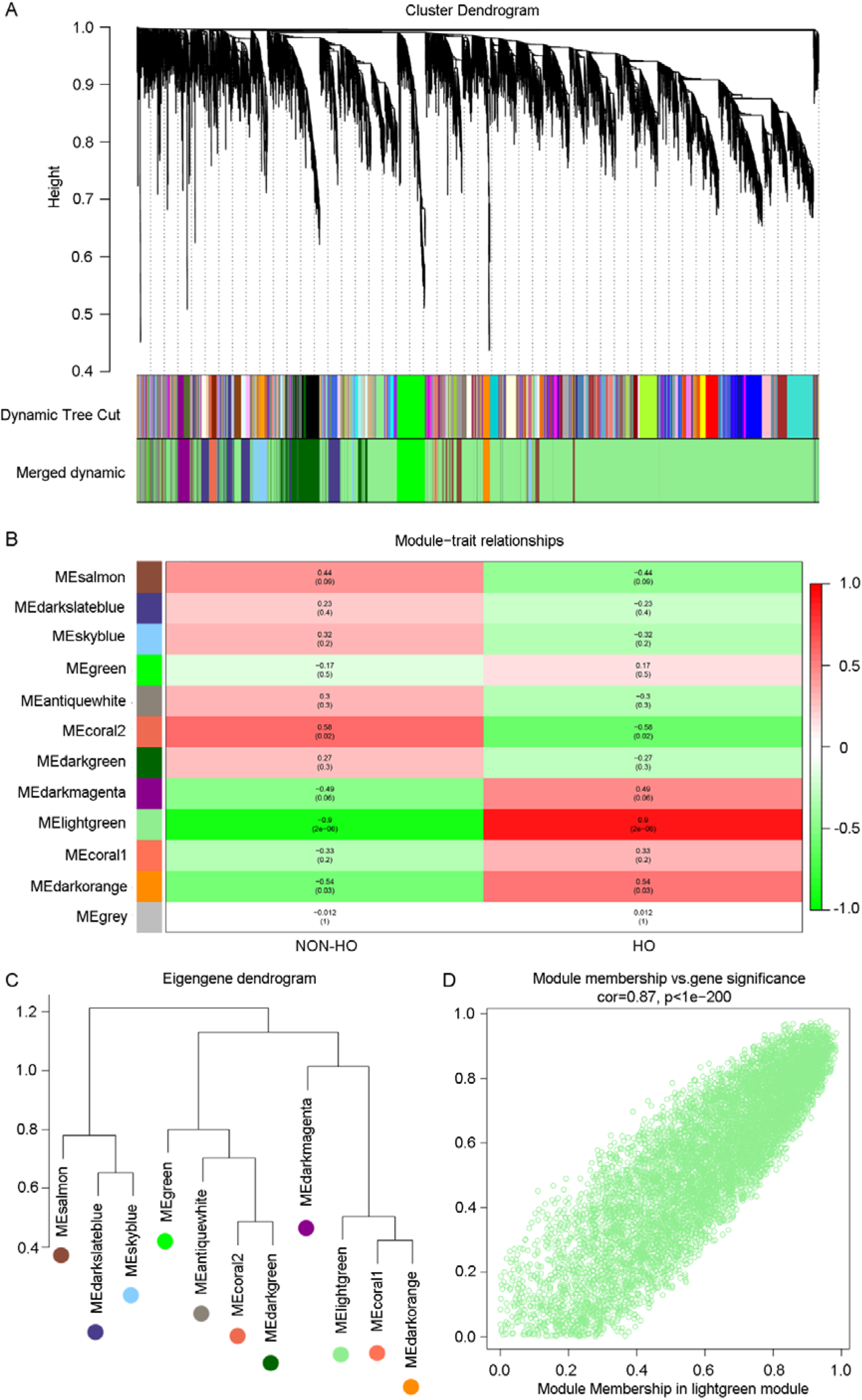
Co-expression network construction and HO-related hub module identification in HO. **A** The tree diagram of each DEG clustered according to different metrics (1-TOM). We established twelve co-expression modules, as displayed in unique colors. **B** Heatmap showing relationship of modules with disease features. In this module, a greater average gene correlation level indicated the closer relationship between module and interested features. Horizontal axis stands for clinical factors, whereas vertical axis indicates modules. Colors from red to green indicates transition from positive into negative correlations. Data within the grid stand for correlation coefficients. The value within brackets is p value of correlation test. **C** Module hierarchical cluster tree. Dendrogram showed characteristic genes within consensus module obtained by WGCNA. **D** Scatter diagram showed module eigengenes in lightgreen module.

### Validation of HO-related hub genes

We employed “limma” package in R language to screen DEGs from GSE94683 dataset on the basis of |log_2_FC| ≥ 2 and *P* < 0.05 as criterions. We screened altogether 1,483 DEGs to be differentially expressed in HO samples compared with NON-HO samples, including 756 with up-regulation and 727 with down-regulation. We later conducted FDR correction by Benjamini and Hochberg approach. The test set GSE126118 dataset was analyzed using the same methodological procedures. The top 5% genes (n=64) from PPI network and top 5% genes (n=413) from co-expression network were overlapped to screened the real hub gene. On the basis of these results, 4 hub genes (SPP1, PTK2, CSF1R, and AGT) associated with HO were screened out from both co-expression networks and PPI networks (Figure 3A). The volcano plot of the DEGs was shown in Figure 3B. Among these genes, SPP1, CSF1R and AGT were upregulated while PTK2 was downregulated. Thus, the 4 genes above were identified as HO related real hub genes to move on to the later analyses.

**Figure 3.**
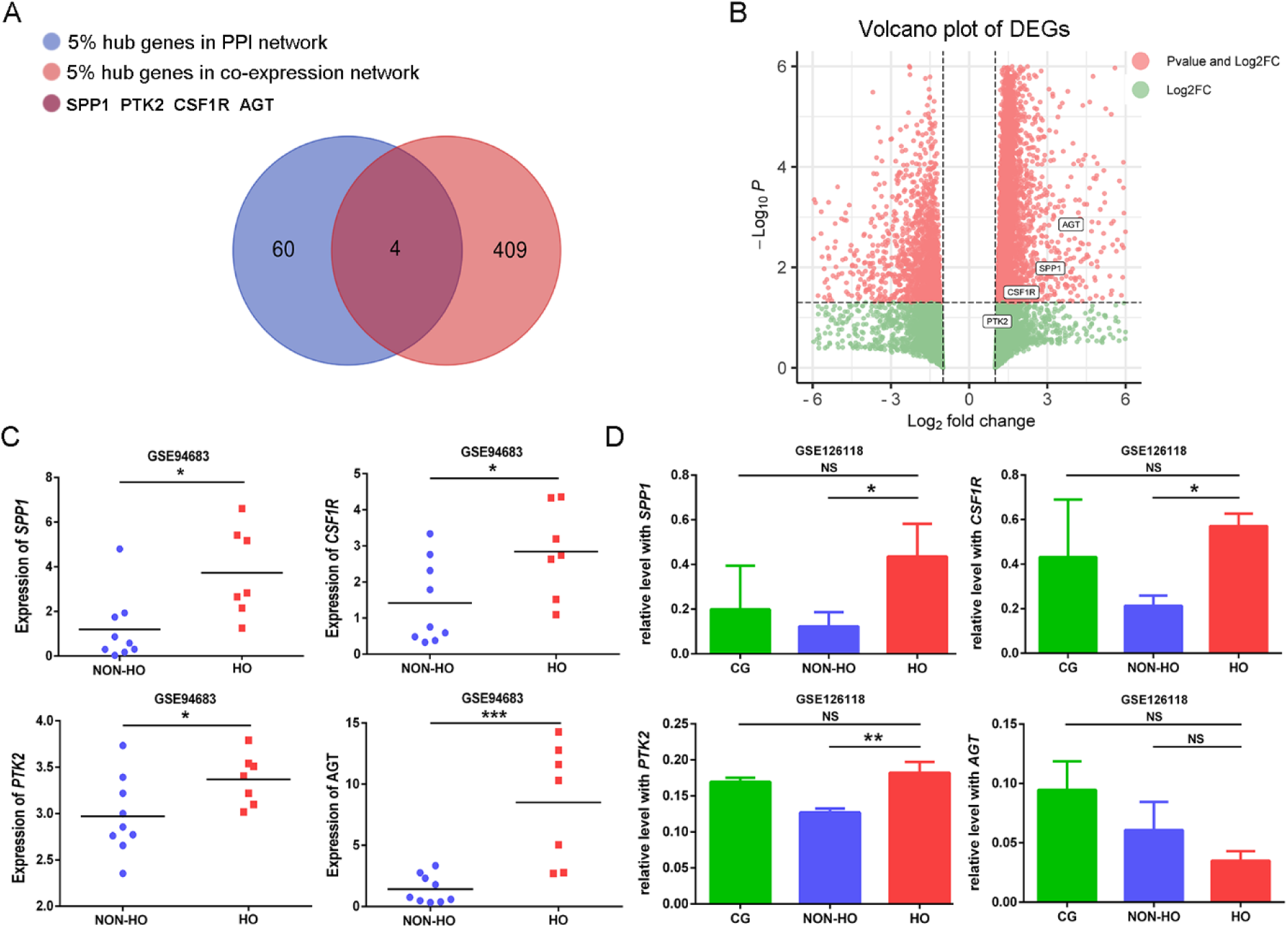
Validation of HO-related hub genes. **A** The volcano plot of DEGs. The thresholds are set as *P* < 0.05 and |log2FC|≥1.0. Red nodes indicate up-regulated genes, while blue ones indicate down-regulated genes. **B** Four hub genes (SPP1, PTK2, CSF1R, and AGT) overlapped between co-expression networks and PPI networks. **C** The validation of hub genes using training set (GSE94683). **D** The validation of hub genes using test set (GSE126118).

To further explore the real hub genes associated with HO, the GSE94683 training set and GSE126118 test set were both utilized to detect the expression levels of SPP1, PTK2, CSF1R and AGT, respectively. Compared with NON-HO samples, each hub gene showed significant up-regulation within HO samples in the training set (Figure 3C) and were excluded AGT from test set (Figure 3D). Results of both datasets were overlapped, three hub genes (SPP1, PTK2 and CSF1R) were found to be significantly altered in HO samples.

### Functional annotations of DEGs

Based on GO functional enrichment analysis, “extracellular space” was identified as one of the most significantly enriched gene sets in HO (Figure 4A). As a result, HO was related to proteinaceous extracellular matrix, extracellular region, extracellular matrix organization, cell adhesion, plasma membrane, positive GTPase activity regulation, negative cell migration regulation, inflammatory response, and heparin binding (Figure 4B). In the meantime, according to KEGG analysis, we found that DEGs showed the most significant enrichment in Pathways in cancer, Cytokine-cytokine receptor interaction, Rap1 signaling pathway, Stem cells pluripotency regulatory pathways, Axon guidance, Steroid biosynthesis, Complement and coagulation cascades, ECM-receptor interaction, and PI3K-Akt pathway (Figure 4B).

**Figure 4.**
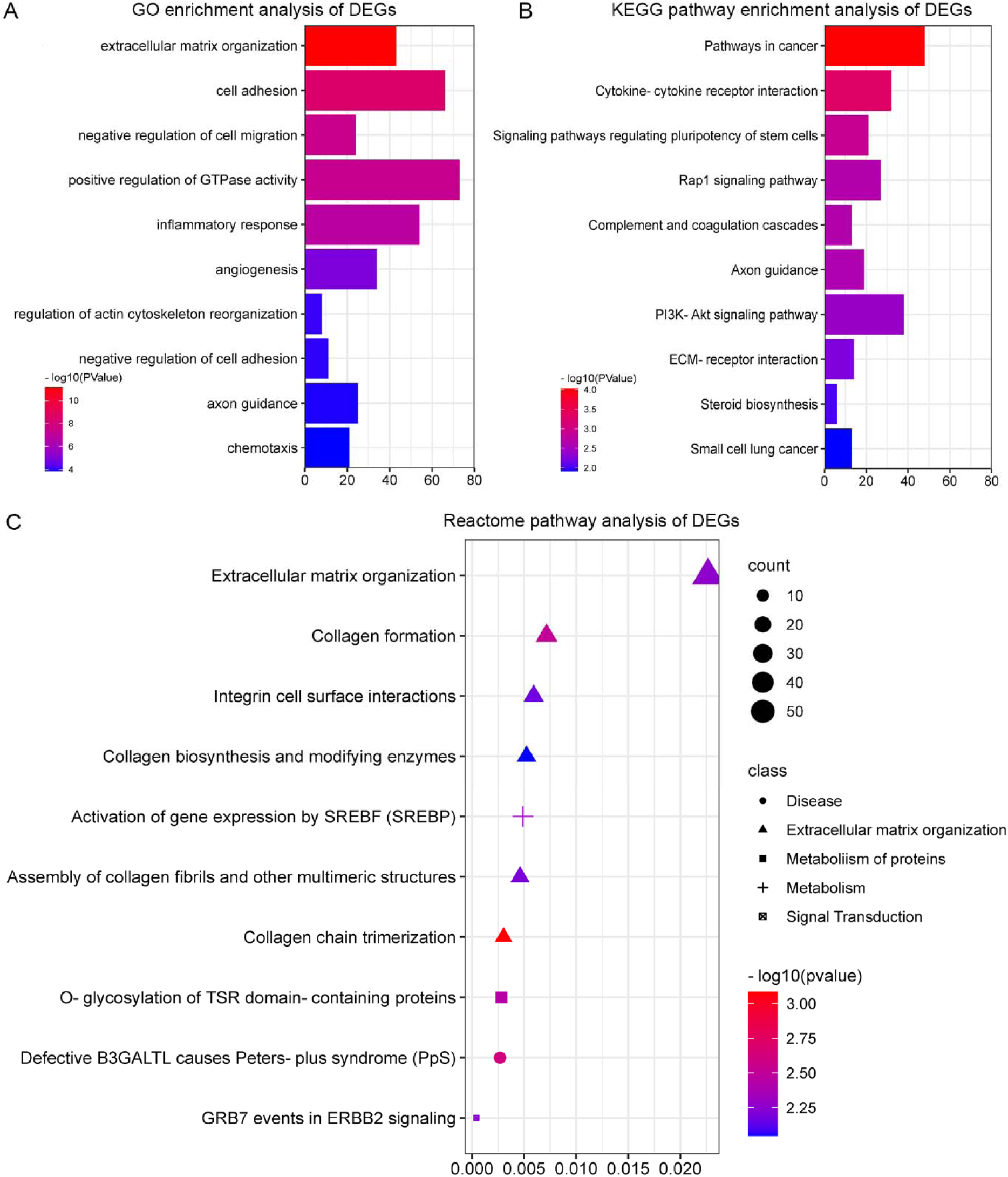
Functional annotation for hub genes. **A** The 10 most significant enriched functional clusters of DEGs. **B** KEGG signal pathway enrichment on DEGs. **C** Bubble diagram shows 10 most significant pathways with Entities FDR, Entities ratio and Entities found (count). X-axis indicates enriched factor, while Y-axis stand for the name of pathway; The color and size of each bubble respectively indicate the -log10 (q value) and the number of DEGs related to the pathway. Gene set participated in HO were significantly enriched in extracellular matrix organization, signal transduction, and metabolism.

Subsequently, Reactome was used as a functional enrichment analysis method to compare the target with those related biological activities. Furthermore, we displayed BPs in bubble chart based on Entities ratio, Entities found, and Entities FDR functions. As a result, HO samples were significantly enriched in Defective B3GALTL causes Peters-plus syndrome (PpS), Collagen chain trimerization, O-glycosylation of TSR domain-containing proteins, Collagen formation, Extracellular matrix organization, Activation of gene expression by SREBF (SREBP), Assembly of collagen fibrils and other multimeric structures, GRB7 events in ERBB2 signaling, Collagen biosynthesis and modifying enzymes, and Integrin cell surface interactions. The top 10 important functional pathways are classified according to entities. From the histogram, it can be observed that these synthesized pathways exert an important part on Signal Transduction, Extracellular matrix organization, Metabolism of protein, Diseases of metabolism, and Metabolism (Figure 4C).

### GSVA and GSEA

In contrast to GO and KEGG functional analyses, GSEA is not conducted by detecting the entire DEG list but instead of validating the functional annotations of the whole genome at transcription level. We conducted GSVA and GSEA for analyzing biological pathways associated with HO. The results of the GSVA showed that HO status was significantly enriched in biosynthesis of unsaturated fatty acids (Figure 5A). The results of GSEA showed that gene sets related to HO were also enriched positively with highest significance in “Proteasome”. The present study also demonstrated that HO samples were enriched in Oxidative phosphorylation, the Protein export, the Lysosome lysosome, the Ribosome ribosome, the Vibrio cholerae infection, the Spliceosome spliceosome, and the Biosynthesis of unsaturated fatty acids (Figure 5B). Top two most significant pathways identified using GSEA were showed in Fig. 5C-D.

**Figure 5.**
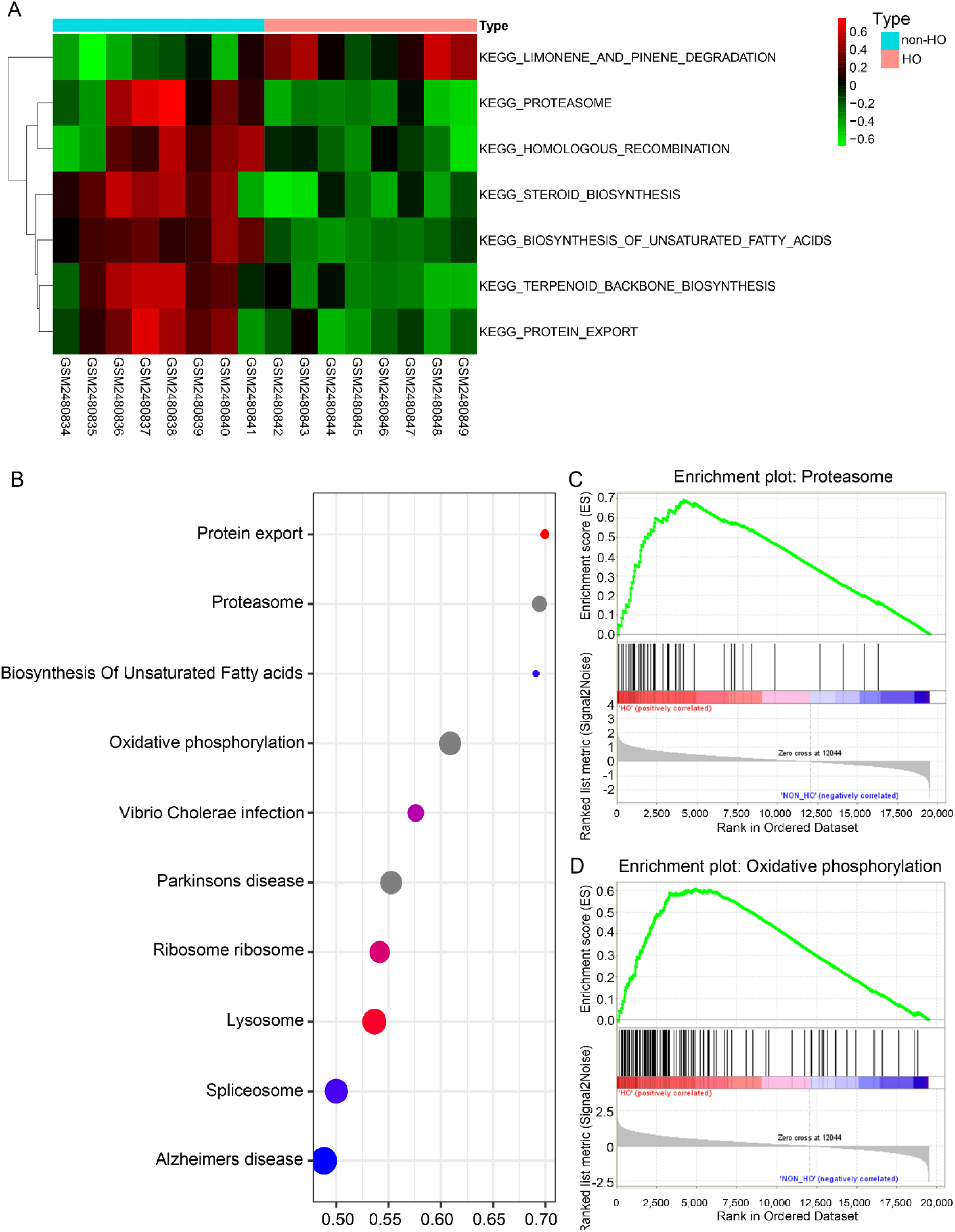
Potential pathway analyses of hub genes. **A** Pathway analyses using GSVA. The heatmap shows HO-relate KEGG gene set enriched using GSVA algorithm. **B** Pathway analyses using GSEA. In the bubble plot, X-axis indicates enriched factor, while Y-axis stand for the name of pathway; The color and size of each bubble respectively indicate the -log10 (q value) and the number of DEGs related to the pathway. **C-D** Top two most significant pathways identified using GSEA. In the heatmap in GSEA results, red and blue stand for up-regulated and down-regulated genes, respectively.

### Evaluation of the relationship between HO and immune infiltrating degrees

We performed CIBERSORT deconvolution for inferring absolute level of immune infiltrate within each sample of HO (Figure 6A). Positive correlation was observed between Type I IFN Reponse and Parainflammation (R2 = 0.83), according to Figure 6B. Instead, Tfh was markedly negatively related to Treg (R2 = −0.78). For better understanding the relation of HO with immune infiltrating degrees, we conducted ssGSEA analysis. 3 TIICs types (T helper cells, neutrophils, and pDCs, and) and 3 immune functions (APC co inhibition, CCR, and MHC class I) showed obvious relation with HO-related risk score of GSE94683 dataset (Figures 6C).

**Figure 6.**
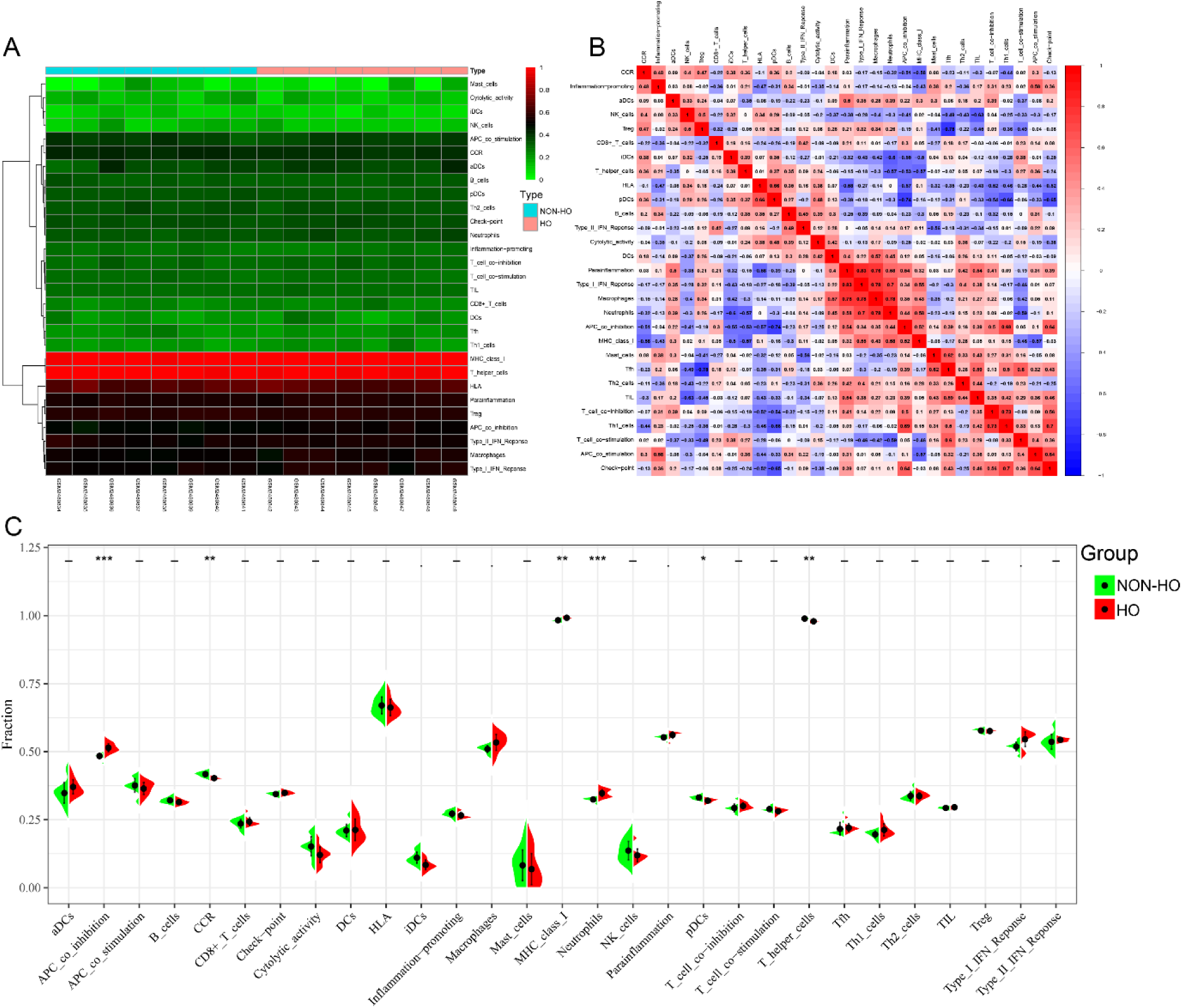
Evaluation on immune infiltration in heterotopic ossification (HO). **A** Heatmaps show highly differentially expressed genes for each cluster of immune cells. The horizontal axis indicates each sample. Samples are sorted by HO and NON-HO. The vertical axis indicates each immune cell. The immune cells with similar expression are visualized through clustering. **B** Heatmap of correlation analysis between various immune cells. The red color reflects a strong association of 2 immune constituents. The blue color reflects the weak association. The color shade represents association strength. **C** Violin plots for different immune cell percentages. The black dot in the middle is the median value. \**P* < 0.05, NS, not significant.

### Identification of potential drugs

The possible molecular compounds or drugs for reversing up-regulated hub genes of HO were predicted by using DGIdb (version 4.2.0). Six drugs or molecular compounds (including calcitonin, gentamicin, tacrolimus, and wortmannin) were found to regulate the expression of SPP1 differentially within the drug–gene interaction network (Figure 7A). Ten molecular compounds or drugs including defactinib were detected to interact with PTK2 (Figure 7B). In addition, we discovered that ten molecular compounds or drugs, like, edicotinib, cabiralizumab, pexidartinib and emactuzumab, interacted with CSF1R (Figure 7C).

**Figure 7.**
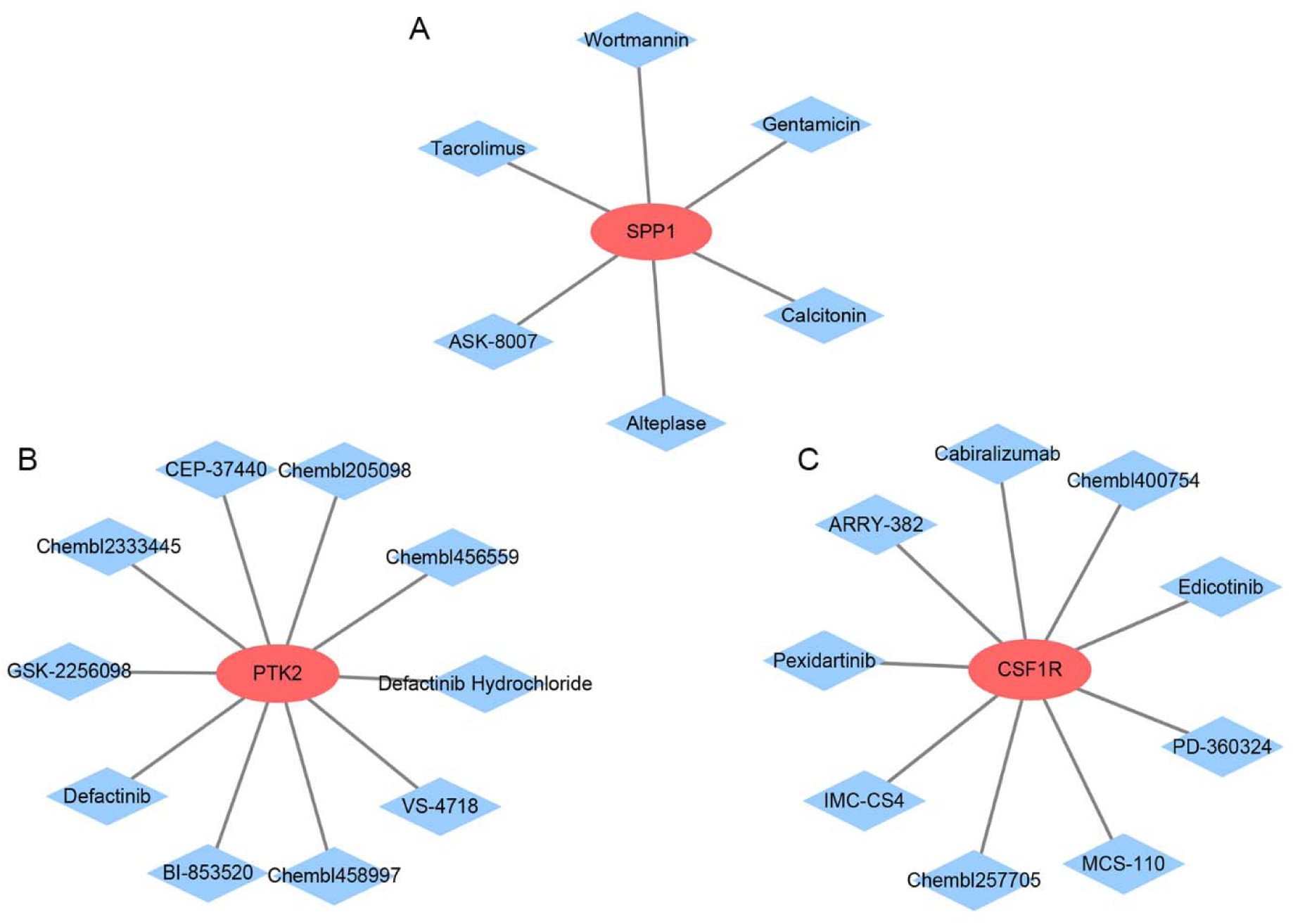
Drug-target interaction network to predict therapeutic potentials. The red ellipse reflects genes, while the light-blue V nodes reflect drugs. **A** SPP1. **B** PTK2. **C** CSF1R.

## Discussion

We identified HO related hub genes by using a comprehensive bioinformatics analysis, including an overlap method that combined WGCNA and PPI network. Meantime, functional enrichment analysis was performed to recognize associated biological pathways. In accordance with our WGCNA findings, we found that lightgreen module in HO phenotype was clinically significant. Subsequently, we identified 4 genes (SPP1, PTK2, CSF1R, and AGT) screened from overlapping PPI and co-expression networks as the real hub genes, which indicated their critical functions in HO genesis and development. Next, to verify those real hub genes in HO, we used both datasets to detect the expression levels of these 4 genes, respectively. Finally, SPP1, PTK2 and CSF1R in both training set and test set revealed significant up-regulation between HO samples and NON-HO samples.

We also annotated the potential enrichment pathways for clarifying DEGs’ biological roles. In the GO analyses, HO-related DEGs is mainly related to proteinaceous extracellular matrix, extracellular region, extracellular matrix organization, and plasma membrane. Consistent with KEGG enrichment analyses, DEGs in HO were mostly enriched in cytokine-cytokine receptor interaction, stem cell pluripotency regulatory pathways, and rap1 signaling pathway. Besides, based on Reactome analysis, HO specimens were closely related to collagen chain trimerization, collagen formation, and O-glycosylation of TSR domain-containing proteins. Subsequently, GSVA and GSEA also demonstrated that the HO related hub genes are most highly enriched in immune responses. Consistent with our study, previous studies confirmed that injury-induced inflammation paramount influences on HO pathogenesis[38, 39]. Moreover, many immune cells and inflammatory molecules also have an essential function in modulating persistent tissue damage response as well as in promoting tissue repair attempts [40]. Inflammatory response is mainly relevant to immune responses related to neutrophils, macrophages and lymphocytes. Previous studies demonstrated that pathological conversion to ossification from calcification after tissue damage is a possible mechanism leading to HO formation [41]. Tissue damage may result in calcium mineral depositing in degenerated tissue. Most calcium is usually effectively removed through phagocytic macrophages; thus, injured tissue can recover.[42]. But calcium mineral can progress into HO when tissue repair fails, and connective tissue cells take an active part in forming collagen matrix [43]. Consistent with these results, more recent studies indicated that macrophage phagocytosis and inhibiting macrophage infiltration in injured tissues enhance HO [43–45]. However, some studies indicated that depletion of macrophages can prevent the formation of HO [46, 47]. Therefore, the role of macrophages in HO needs further exploration. In addition, immune cells, like resident cells, lymphocytes, neutrophils and macrophages, produce various chemical molecules, such as purines, lipids, histamine, protons, bradykinin, serotonin, chemokines, cytokines and nerve growth factors in inflammation [48]. Chemokines can influence osteogenic differentiation, which are also upregulated by injury or inflammation [49]. Collectively, previously published results showed that heterotopic ossification displayed intricate biological processes, which involves the coordination of immune regulation signals and inflammatory response after soft tissue injury.

To further validate the HO related hub genes, we acquired gene expression patterns in GEO. As expected, those four genes collected from the above database, including SPP1, PTK2 and CSF1R, were highly expressed in HO samples. These suggested that three hub genes above were significantly related to HO. Secreted phosphoprotein-1 (SPP1), which is referred to as osteopontin (OPN) as well, can be produced to bind to hydroxyapatite and participate in osteoclast attaching into mineralized bone matrix. SPP1 is reported to show positive relation with HO while its up-regulation in posterior longitudinal ligament cells [50]. Besides, a study of hypophosphatemic osteosclerosis found that alterations in SPP1 were associated with HO [51]. Another study of ossification of the ligamentum flavum found that SPP1 was involved in the process of miR-615-3p unfavorably regulating ossification in lumbar ligamentum flavum cells of human beings [52]. However, a study from Crowgey EL et al. [53] suggested SPP1 down-regulation within HO+ samples relative to HO-samples, consistent with a previous study. PTK2 (coding as FAK) is the cytoplasmic protein tyrosine kinase that shows concentration within focal adhesions formed between cells that grow with ECM components. In accordance with Huber AK and colleagues [54] discovered that PTK2 up-regulation was related to HO, while pharmacologic PTK2 inhibition or genetic deletion of PTK2 within MPC lineage cells alleviated HO. The colony stimulating factor-1 receptor (CSF-1R) accounts for an important regulating factor for the occurrence and maintenance of tissue macrophages [55]. To the best of our knowledge, the present work suggested CSF1R up-regulation within HO samples relative to NON-HO samples. But, there are less reports related to the effects of CSF1R on HO due to it is a relatively new molecules at present. Consequently, more studies are needed for exploring CSF1R’s effect on HO. To sum up, these genes may provide us new orientation in experimental research and clinical study. Further research is required to fully explore their roles in HO.

Additionally, DEGs and infiltrating degrees of immune cells were analyzed between low and high expression groups of hub genes. As a result, immune cells infiltrates increased within one immune cells (neutrophils) and two immune associated features (APC co inhibition, and MHC class ), while immune cells (pDCs, and T helper cells) and one immune associated feature (CCR) were decreased. Such results conform to current findings. Prior works have found that APC co inhibition and MHC class could contribute to the progression of osteogenesis differentiation and to the activation of immune responses in bone marrow mesenchymal stem cells [56, 57] , which is consistent with our findings. Recent study has shown that traumatic injury can lead to initiation of a cascade of neurogenic inflammation, thus leading to the recruitment of neutrophils to the injury site [58]. Moreover, neutrophil-mediated immune cell-based pathways play a key part in burn-mediated HO [59]. Based on such findings, immune response possibly has a critical function in HO development. However, associations among MHC class , APC co-inhibition and HO was not found in the previous study. Therefore, exact functions of MHC class and APC co-inhibition in this process of HO needs to be further studied.

To predict therapeutic drugs possibly reversing up-regulation of OH-associated hub genes, DGIdb database was performed to recognize the potential effective treatment of OH and OH-related complication. Viana R et al. [60] used calcitonin to treat recalcitrant phantom limb pain (PLP) with concurrent heterotopic ossification. As a result, calcitonin contributed to managing PLP and HO in several disease occurrence and maintenance stages. An earlier study about trauma-induced heterotopic ossification showed that gentamicin can decrease the infection in traumatic arthritis patients [61]. However, other drugs, including tacrolimus, wortmannin, defactinib, edicotinib, cabiralizumab, pexidartinib and emactuzumab, have not been reported to be used for HO treatment to date. Tacrolimus has previously suggested to cause osteoporosis through up-regulating osteoclast-differentiation factor expression or suppressing osteoblast differentiation through restraining Runx2 expression and calcineurin activity [62, 63]. As a kind of fungal metabolite, Wortmannin specifically inhibits the phosphatidylinositol 3-kinase (PI3K) family including ataxia telangiectasia mutated kinase and double-stranded DNA dependent protein kinase [64]. Defactinib is a kind of small-molecular focal adhesion kinase inhibitor with oral availability, which can suppress OA through inhibiting positive feedback loop between MSCs and H-type vessels within subchondral bone. Edicotinib accounts for a colony-stimulating factor-1 (CSF-1) receptor inhibitor with selectivity and oral availability, which is currently entering the phase IIA clinical trial against rheumatoid arthritis (RA) [65]. Cabiralizumab is a humanized monoclonal antibody that specifically targets colony stimulating factor-1 receptor (CSF1R), and it belongs to CSF1/PDGF tyrosine-protein kinase family [66]. As CSF1R’s receptor tyrosine kinase inhibitor Pexidartinib has anti-cancer effect [67]. Emactuzumab represents the prominent anti-CSF-1R antibody, which can specifically target or eliminate tumour-associated macrophages (TAMs) [68]. More studies are warranted to explore the effects of those aforementioned molecular compounds and drugs on HO along with the related diseases.

Nonetheless, certain limitations should still be noted. First, although our identify of HO-related novel hub genes and biological pathways, there are finitudes of available microarray data and inherent bias of enrichment analysis to these data. Second, the quality of the data cannot be appraised, due to gene expression profiles were obtained from public database. Third, tissue samples from Homo sapiens in training set differ from Mus musculus in test set, which may generate different target genes between two organisms after HO. Consequently, further investigations are needed in the case of sample updates in the database. Fourth, it is necessary to further verify the results by laboratory experiment. Last, more tissue samples are required herein to be repeated increasing the reliability of results. Therefore, further researches are needed for confirming the effects of these core genes as well as the biological pathways on HO genesis and progression.

## Conclusion

In conclusion, we carried out comprehensive integrative profiling and bioinformatics analysis to identify 3 HO-related hub genes of high expression and to predict potential therapeutic drugs. The enrichment biological pathways conformed to the existing understanding of disease pathogenesis. Research on such hypothesis can shed more lights on HO-related biological pathways, and determine multiple candidate regulatory factors as intervention targets. However, more research is warranted for confirming the relation of hub gene functions with immune responses in HO development.

## Abbreviations

HO: Heterotopic ossification
GEO: Gene Expression Omnibus
DEGs: Differentially expressed genes
WGCNA: Weighted gene co-expression network analysis
PPI: protein-protein interaction
GO: Gene Ontology
KEGG: Kyoto Encyclopedia of Genes and Genomes
GSVA: Gene set variation analysis
GSEA: Gene set enrichment analysis
MM: Module membership
GS: Gene significance
TOM: Topological overlap matrix
DGIdb: Drug-Gene Interaction database
STRING: Search Tool for the Retrieval of Interacting Genes
FC: fold change
FDR: False discovery rate
SPP1: secreted phosphoprotein 1
PTK2: protein tyrosine kinase 2
CSF1R: colony-stimulating factor 1 receptor
AGT: angiotensinogen

## Acknowledgements

Our thanks should go to GEO database for offering the platforms and uploading the useful datasets.

## Authors’ contributions

MLY designed data analysis and wrote the manuscript. YPS, SCS, BY LY acquired and analysed the data. BY, LY, LXL edited the manuscript. BY, LY, LXL contributed to critical revision. All authors have read and approved the manuscript.

## Availability of data and materials

All supporting data can be provided upon request to the authors.

## Declarations

### Ethics approval and consent to participate

This article does not contain any studies with human participants or animals performed by any of the authors.

### Consent for publication

All the authors approved the manuscript.

